# Autoantibodies from recovered Guillain-Barré syndrome patients exhibit altered effector functions that prevent subsequent neuron degeneration

**DOI:** 10.64898/2026.08.31.748457

**Authors:** Ashley M. Rogers, Jessica L. McAlpine, Xu Yang, Israt Jahan, Nowshin Papri, Shoma Hayat, Stephanie Archer-Hartmann, Meagan Shinn, Parastoo Azadi, Nadja Zeltner, Zhahirul Islam, Christine M. Szymanski

**Affiliations:** Department of Microbiology, University of Georgia, Cedar Street Building C, 120 Cedar St., Athens, GA 30602, USA; Complex Carbohydrate Research Center, University of Georgia, 315 Riverbend Drive, Athens, GA 30602, USA; Center for Molecular Medicine, University of Georgia, 325 Riverbend Road, Athens, GA 30602, USA; Department of Biochemistry and Molecular Biology, University of Georgia, Davison Life Sciences Complex, 120 East Green St., Athens, GA 30602, USA; Gut-Brain Axis Laboratory, Infectious Diseases Division, icddr,b, 68, Shaheed Tajuddin Ahmed Sarani Mohakhali, Dhaka 1212, Bangladesh; Department of Cellular Biology, University of Georgia, Biological Sciences Building, 1000 Cedar St., Athens, GA 30602, USA

## Abstract

Guillain-Barré syndrome (GBS) is an autoimmune polyneuropathy that is the leading cause of nonpoliovirus-associated acute flaccid paralysis worldwide. In most cases, GBS occurs following an infection, most commonly *Campylobacter jejuni*, through the induction of antibodies recognizing bacterial ganglioside-mimicking lipooligosaccharides that cross-react with human neuronal gangliosides. This post-infectious autoimmune cascade causes neuropathy, from which patients can recover as their anti-ganglioside antibody titers diminish and immunostimulation decreases. In this study, we find 10% of clinically confirmed GBS patients maintain high titers of circulating anti-ganglioside antibodies more than one decade after recovery. These antibodies no longer cause neuropathy compared to acute sera from the same patients using a human pluripotent stem cell-derived sensory neuron model with human complement. We found both IgG subclass and glycoform differences between paired acute and recovered GBS patient sera, including anti-inflammatory modifications on isolated anti-GM1 ganglioside antibodies. Together, these data suggest that patients with GBS select for non-pathogenic variants of autoantibodies that are no longer capable of damaging their neurons, but may still protect against *C. jejuni* infection.

## Introduction

Guillain-Barré syndrome (GBS) is an autoimmune disease that causes peripheral neuropathy and is the leading form of nonpoliovirus-associated acute flaccid paralysis^1^. GBS typically presents with weakness, symmetrical limb paralysis, and numbness in the extremities; mortality can occur once paralysis reaches the lungs and is left untreated^2^. The rate of incidence for GBS is 1:100,000 worldwide, but populations in low- and middle-income countries (LMICs) are at a 2-fold greater risk of developing GBS^3^. Approximately 40-70% of patients develop GBS following an infection, most commonly with *Campylobacter jejuni*, a leading cause of bacteria-associated enteritis worldwide^4,5^. Infection with *C. jejuni* can lead to the generation of cross-reactive antibodies against the bacterial ganglioside-mimicking lipooligosaccharides (LOS) that also recognize human gangliosides on neurons^6^. In most individuals, peripheral tolerance removes self-reactive B cells which often prevents autoantibody production and subsequent GBS development. However, in approximately 1 in 1,000 *C. jejuni* infections, autoantibodies develop against the LOS which cross-react to the gangliosides found on sensory and motor neurons, triggering the paralysis observed in GBS patients^5,7,8^. Furthermore, in *C. jejuni* associated GBS cases, at least 73% of the isolated *C. jejuni* are ganglioside-mimicking^9^. The high titers of anti-ganglioside antibodies can then bind to human neuronal membranes and initiate a cascade of events resulting in neuron inflammation and degeneration. These events are comprised of complement deposition leading to reactive oxygen species (ROS) release, demyelination, lesions at the nodes and Wallerian-like degeneration of the neuron^10–12^.

GBS comprises several clinical and electrophysiological subtypes, including acute motor axonal neuropathy (AMAN), acute inflammatory demyelinating polyradiculoneuropathy (AIDP), acute motor sensory axonal neuropathy (AMSAN), and Miller Fisher syndrome (MFS) resulting from damage to different types of neurons which vary in their expression of ganglioside type and abundance^13^. For example, AMAN is most commonly associated with antecedent infections and anti-GM1 antibodies, which can contribute to complement-mediated degeneration of motor neurons. However, some *Campylobacter*-related GBS cases can also present as the demyelinating (AIDP) type^14,15^.

Patients with prolonged high titers of anti-ganglioside antibodies have been associated with more severe outcomes and paralysis^2,16^. Therefore, the main treatment for GBS involves the removal of these pathogenic antibodies using plasma exchange or intravenous immunoglobulin (IVIg)^17^. However, a substantial portion of patients (40% of GBS) experience delayed or incomplete improvement after these treatments suggesting a need for more effective therapeutics^18^.

Specific antibody glycoforms are one promising avenue for improved GBS therapeutics. Antibody modification by glycosylation of asparagine 297 on the Fc portion of IgG profoundly impacts inflammation by modulating binding affinities for physiologically relevant targets. Patients with autoimmune diseases frequently tend to have low IgG N-glycan galactosylation, sialylation, and fucosylation^19^. The effects of IgG glycosylation are reflected in how different IgG glycoforms impact complement-dependent cytotoxicity (CDC). For instance, increased galactosylation can increase inflammation by permitting better antibody positioning and binding to C1q^20^. Conversely, sialylation of the N-glycan has poorer affinity for C1q leading to less CDC^21^. Indeed, patients who have higher levels of sialylation tend to be enter remission sooner in related diseases like chronic inflammatory demyelinating polyneuropathy (CIDP)^21^. The IgG glycoforms associated with GBS are more variable than those found in CIDP; 24.2 to 71.4% of IgG is galactosylated, whereas 5.9 to 26.0% is sialylated^22^. Furthermore, fucosylation of the N-glycan renders IgG anti-inflammatory due to decreased affinity to FcγRIIIa expressed on phagocytes and decreased complement activation^23,24^. Changes in fucosylation have not previously been reported in GBS, but afucosylation is more common in autoimmune diseases compared to higher rates of fucosylation in healthy populations^25–27^.

In this study, we compare IgG subclasses and glycoforms from the same donor during acute GBS and more than one decade later following their recovery. We report here that just over 10% of GBS patients in our population retain circulating anti-GM1 IgG antibodies following resolution of paralysis. To examine IgG pathogenicity, we created a novel GBS *in vitro* neuropathy model using human pluripotent stem cell (hPSC)-derived sensory neurons treated with commercial and patient antibodies. Here, we recapitulate complement-mediated neuron-induced damage from acute phase IgG but, not IgG post-GBS. To determine why these autoantibodies do not cause neuropathy, we characterized their IgG glycoforms and subclass composition. Remarkably, we were also able to isolate enough antigen specific anti-GM1 antibodies from recovered GBS patients to compare those to total IgG in the same samples. Furthermore, in some patients, we detected cross-reactive antibodies to our *C. jejuni* laboratory isolates with GM1-mimicking LOS. The results described here lead to a new paradigm suggesting that changes in antibody post-translational modifications leads to less neuronal complement-mediated damage while retaining the ability to reduce *C. jejuni* growth, providing an explanation into why these autoantibodies remain in circulation despite their potential pathogenicity.

## Results

### Anti-ganglioside antibodies are found in recovered GBS patients

Since anti-GM1 antibodies are among the most common anti-ganglioside antibodies detected in GBS patients, anti-GM1 ganglioside antibody titers were assessed in sera from recovered GBS patients, recovered *C. jejuni* enteritis patients, and healthy controls^28^. Out of 100 recovered GBS donors, 10 still had high circulating anti-GM1 ganglioside antibodies, and at significantly greater titers than the healthy and enteritis cohorts (Fig. 1a). Of note, out of 100 healthy donors, there were three people who also showed positive anti-GM1 titers (Fig. 1a), and none of those donors were from the same household as the GBS donors (Table 1). There were no detectable anti-GM1 antibodies from the 26 donors with culture-verified *C. jejuni* enteritis (Fig. 1a). To investigate whether the anti-GM1 antibody positive GBS patients exhibited elevated reactivity to other gangliosides, sera were assessed for presence of circulating antibodies against GA1, GM2, GD1a, GD1b and GQ1b. Of these, only GA1, GD1a, and GD1b were detected in a small subset of this group (Fig. 1b). One former GBS patient (G19) had a high antibody titer against GA1, GD1a, and GD1b ganglioside, while high antibody titers against GD1a and GD1b were also found in a healthy donor (H10) (Fig. 1b). To determine if the prevalence of anti-GM1 is correlated with GBS disease severity, we compared stored sera from 22 patients obtained at the time of GBS diagnosis, referred to as “acute” and compared these to sera collected after GBS recovery from the same patients. Four donors showed high levels of anti-GM1 antibodies in both their acute and recovered sera: G6, G8, G19, and G38 (Fig. 1c). Furthermore, the presence of anti-GM1 antibodies in the recovered GBS patients did not correlate with their previous time of diagnosis or disease severity (Table 1). These data indicate that some recovered GBS patients still have circulating anti-GM1 antibodies although they do not exhibit typical GBS symptoms.

**Fig. 1:**
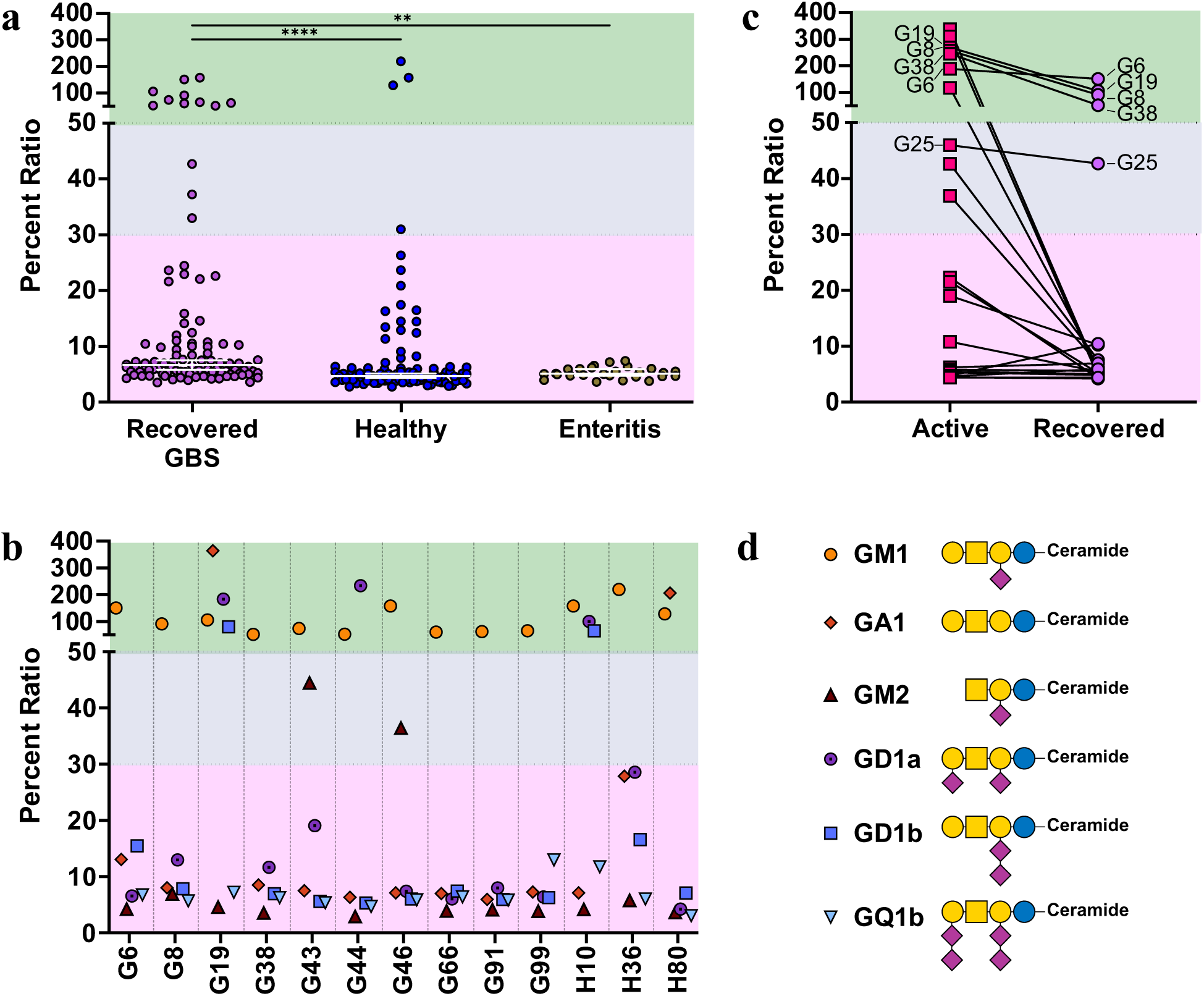
Circulating anti-ganglioside antibodies are present in recovered GBS patients. **a** Comparison of anti-GM1 antibody titers between recovered GBS (*n* = 100), healthy (*n* = 100), and *C. jejuni* enteritis (*n* = 26) groups represented as the percentage of the ratio between the signal intensity of the sample to the calibrator (provided by the manufacturer). **b** Anti-ganglioside antibody percent ratios from donors with anti-GM1 antibodies (*n* = 12). Patient ID starting with a “G” and “H” indicate GBS and healthy donors, respectively. **c** Lines between acute (*n* = 22) and recovered (*n* = 22) groups connect sera collected from the same individual. **d** Illustration of the gangliosides tested. Green highlighted section on the graphs indicate the range where donors are positive for indicated ganglioside antibodies, grey highlighted sections indicate ambiguous presence in binding (as stated by manufacturer), and pink sections indicate negative for anti-ganglioside antibodies. Each data point represents the average of duplicate measurements. Symbol nomenclature for glycans (SNFG)^57^: purple diamond, sialic acid (Neu5Ac); yellow circle, galactose (Gal); yellow square, N-acetylgalactosamine (GalNAc); blue circle, glucose (Glc). Bars in black show the median of each group with a 95% confidence interval (CI). ** *p*-value ≤ 0.01, **** *p*-value ≤ 0.0001 determined by Kruskal-Wallis test.

**Table 1:** GBS patient history. AMAN: Acute motor axonal neuropathy. AMSAN: Acute motor and sensory axonal neuropathy. AIDP: acute inflammatory demyelinating polyradiculoneuropathy. N/A: not applicable. IVIg: intravenous immunoglobulin. SVPE: small volume plasma exchange.

| Patient ID | Diagnosis | GBS diagnosis year | GBS disability Score during GBS | Treatment type | Current Symptoms | Age at enrollment | Household control ID | anti-GM1 Abs |
| --- | --- | --- | --- | --- | --- | --- | --- | --- |
| G6 | AMAN | 2011 | 4 | N/A | Pain and weakness | 48 | H2 | + |
| G8 | AMAN | 2010 | 2 | IVIg | Pain | 18 | N/A | + |
| G19 | AMAN | 2010 | 2 | N/A | None | 48 | N/A | + |
| G38 | AMSAN | 2001, 2011 | 4 | N/A | Weakness, pain | 42 | N/A | + |
| G43 | Unclassified | 2011 | 5 | N/A | Autonomic dysfunction, weakness | 42 | N/A | + |
| G44 | AIDP | 2011 | 4 | N/A | Tremor | 54 | N/A | + |
| G46 | AMSAN | 2010 | 4 | N/A | Sensory deficits, weakness, pain, paresthesia | 37 | H30 | + |
| G66 | AMAN | 2015 | 4 | plasmapheresis | Foot drop, pain | 61 | N/A | + |
| G91 | AMAN | 2014 | 2 | N/A | None | 36 | N/A | + |
| G99 | AMAN | 2012 | 4 | plasma exchange (SVPE) | Pain | 46 | N/A | + |
| G14 | AMAN | 2010 | 4 | N/A | Muscle pain, sensory deficit | 39 | N/A | – |
| G15 | AMAN | 2010 | 4 | N/A | Resting tremor.<br>Fatigue | 22 | H6 | — |
| G26 | AMAN | 2014 | 4 | N/A | Tremor,<br>Fatigue,<br>Palpitation,<br>Poor memory,<br>Asthma | 30 | N/A | — |
| G29 | AIDP | 2015 | 3 | IVIg | Weakness | 46 | N/A | — |
| G32 | AMAN | 2010 | 2 | N/A | pain, resting tremor,<br>breathlessness | 19 | H18 | — |
| G45 | AMAN | 2011 | 4 | plasmapheresis | Weakness,<br>tingling sensation,<br>footdrop,<br>tremor,<br>twitching | 35 | N/A | — |
| G54 | AMAN | 2011 | 4 | N/A | weakness,<br>fatigue,<br>tremor | 22 | H45 | — |
| G60 | AMAN | 2013 | 3 | N/A | Jerking movement | 33 | N/A | — |
| G68 | AMAN | 2011 | 4 | N/A | Tingling sensation | 55 | N/A | — |
| G72 | AMAN | 2012 | 3 | N/A | None | 24 | N/A | — |

### hPSC-derived sensory neurons (SN) model of GBS neuropathy

Many patients in this study reported sensory deficits resulting from damage to sensory neurons (Table 1). To investigate whether the circulating anti-GM1 antibodies in recovered GBS patients could cause neuropathy, we developed a novel GBS model using hPSC-derived SNs. SN identity was confirmed by immunofluorescence (IF) which revealed that SNs co-expressed the peripheral neuron cytoskeleton protein peripherin (PRPH) and the SN-specific transcription factor BRN3A (Supplementary Fig. 1a). SNs were treated with commercial human complement together with either IgG purified from acute sera taken from GBS patients or sera collected from the same patients one decade after they had recovered (Fig. 2a). IF was used to confirm that the antibodies bind to the neurons (Supplementary Fig. 1b). To assess if IgG from patients with acute GBS would cause more neuronal degeneration than IgG from the recovered patients, we used Fluoro-Jade C (FJC) as a marker for neuron degeneration (Fig. 2b-c) ^29,30^. Antibodies were considered neuropathic if the FJC signal that colocalized with propidium iodide, a marker for Nissl bodies, was significantly higher than the FJC signal from healthy IgG (hIgG). As proof of concept, we observed a significant increase in neuron degeneration when neurons were treated with commercial anti-GM1 and anti-GA1 antibodies. The FJC data indicate that the IgG antibodies purified from acute GBS individuals were more destructive than IgG collected after recovery (Fig. 2b-c). All three paired samples showed the same trend although the purified IgG from one patient (G6) did not significantly differ between acute and recovered with the number of samples tested (*p* = 0.0646). In addition to FJC, we measured ROS levels after treatment with the purified IgG antibodies. Again, we see a clear trend suggesting the antibodies from recovered donors are less pathogenic than their acute counterparts, despite these results not reaching statistical significance (Supplementary Fig. 1c-d). Based on the FJC data, which is a more sensitive and specific marker for neuropathy than ROS, these results indicate that anti-GM1 antibodies from recovered GBS patients do not cause neuropathy-like phenotypes while anti-GM1 antibodies from acute GBS can in a novel hPSC-derived sensory neuron model^31^.

**Fig. 2:**
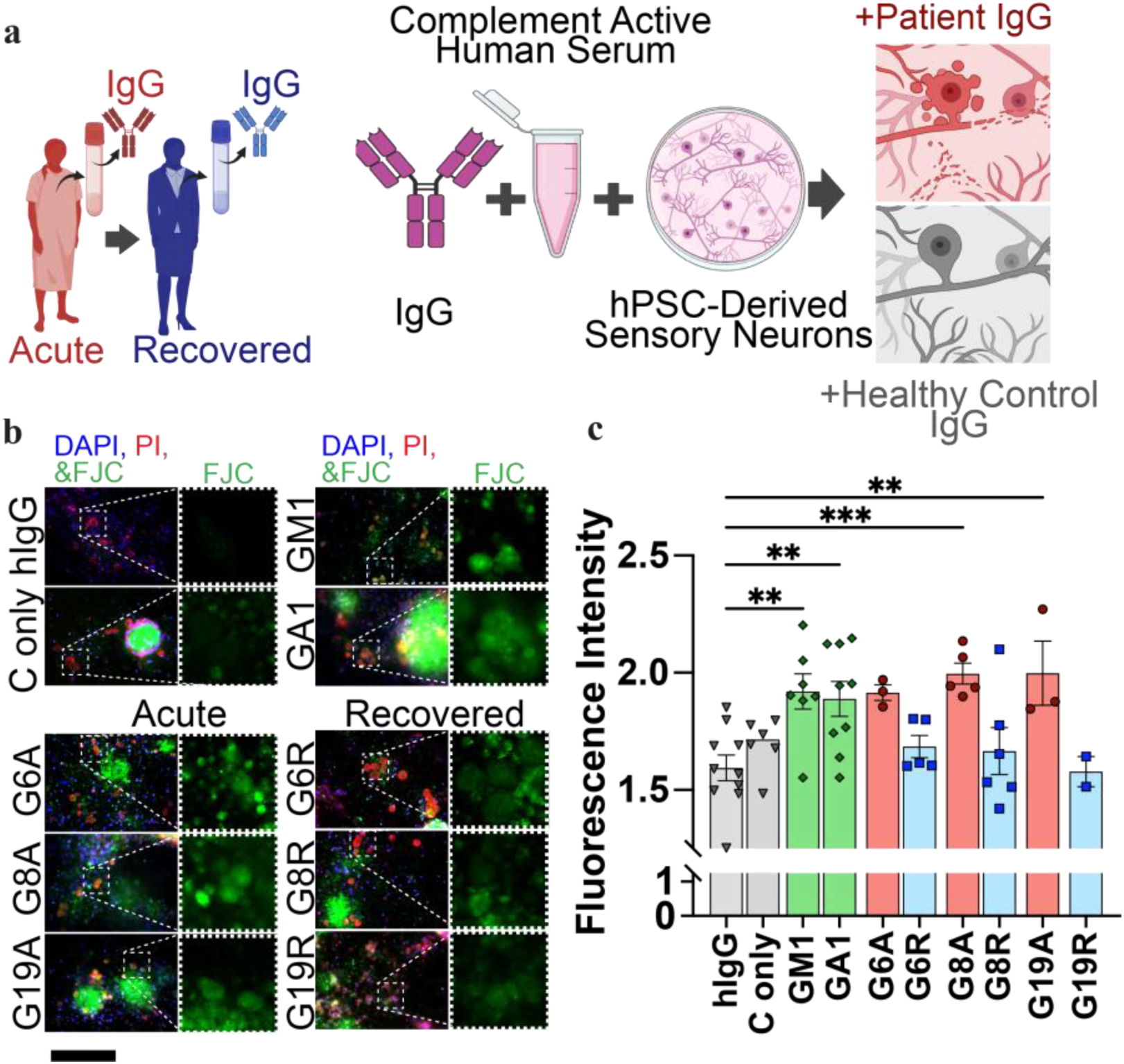
Sensory neuron (SN) degeneration after treatment with IgG. **a** Schematic of SN treatment comparing IgG purified from the same individual at two different time points of patient disease progression: acute (in red) and recovered (in blue). **b** Neuron degeneration was analyzed with Fluoro-Jade C (FJC; green) staining. SN cell bodies were identified using propidium iodide to visualize Nissl bodies (PI, red) and nuclear counterstain DAPI (blue). Magnified insets (dashed white lines) highlight regions with the presence of PI+ cell bodies for closer inspection of FJC signal. Scale bar: 100 µm. **c** Quantification of FJC. Patient samples (red, acute; blue, recovered) were compared to healthy human IgG (hIgG) (grey) by one-way ANOVA. Bars in red indicate IgG from acute GBS timepoint and bars in blue indicate IgG from recovered GBS timepoint. Complement alone (C only) (grey) included as negative control. Commercial anti-GM1 (green) and anti-asialoGM1 (GA1) (green) included to show damage induced by specific anti-ganglioside antibodies. Each dot represents the average of technical duplicates from one biological replicate. Biological replicates are defined as separate SN differentiations from hPSCs, each treated on differentiation day 75. Lines and bars represent mean and standard error of mean. ** *p*-value ≤ 0.01, *** *p*-value ≤ 0.001 determined by one-way ANOVA.

### IgG N-glycan changes in recovered GBS patients

Previous literature indicates that the severity of GBS neuropathy is dependent on the levels of anti-ganglioside antibodies^2,16^. However, recovered GBS donors with high anti-GM1 antibody titers are not showing signs of GBS and their purified IgG showed significantly less FJC and ROS neuron damage compared to the anti-GM1 and anti-GA antibody controls. To investigate why antibodies from recovered GBS patients do not cause neuron damage despite their ability to bind to autoantigens, we examined N-glycosylation of these antibodies on Asn297 since it is known that this modification impacts activation of the classical complement pathway. We obtained this information by glycopeptide compositional data profiling the IgG Fc glycome from GBS patient sera with high anti-GM1 titers and observed several statistically significant differences (Fig. 3a). When examining IgG1 subclass antibodies, IgG from patient G6 recovered sera showed less fucosylation per glycan, and less sialylation normalized to galactose following disease resolution, as compared to IgG collected during acute illness. The IgG2/3a subclass antibodies showed an increase in galactosylation, bisecting GlcNAcylation, and sialylation per galactose compared to IgG collected during acute disease. Patient G6 also showed a decrease in sialylation per galactose in IgG3b/4 compared to their IgG during acute disease. IgG from patient G8 did not exhibit any differences in glycosylation among total antibodies within the IgG1 subclass, but their IgG2/3a showed a similar increase, like patient G6, in galactosylation modifications and an increase in global sialylation. The glycome of IgG3b/4 from patient G8 showed a dramatic decrease in galactosylation and sialylation. Also, the subclass composition showed a significant decrease in IgG1 from patient G8 (Fig. 3b). Patient G19 had a decrease is galactosylation in each subclass. Patient G38, who had two recurrences of GBS following their initial diagnosis, did not exhibit a difference in IgG subclass or glycome comparing the original acute serum sample to the recent sampling (Supplementary Fig. 2). This suggests an IgG proinflammatory glycosylation profile that we are planning to test in our sensory neuron model (in addition to expanding our patient number of recurrent GBS cases). Overall, all three of the recovered GBS donors with high titers of anti-GM1 antibodies showed changes in total IgG Fc glycoforms consistent with a less proinflammatory profile.

**Fig. 3:**
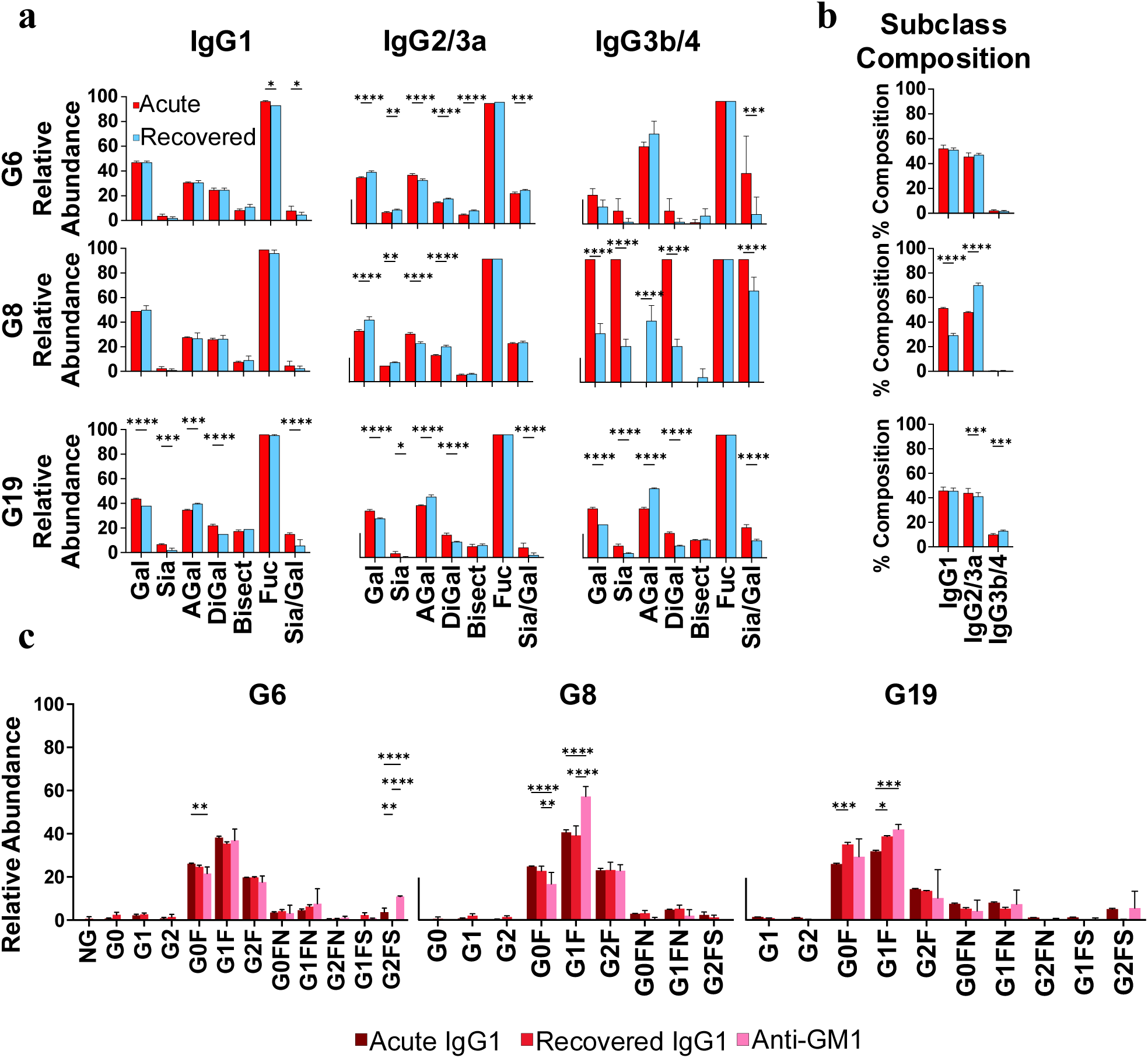
Liquid chromatography-mass spectrometry analyses of GBS patient sera during acute and recovered states. **a** Glycosylation of the Fc portion of IgG separated by subclasses IgG1, IgG2/3a, and IgG3b/4. Relative abundance refers to normalized peak area detected for each glycoform. **b** Percent composition of each IgG subclass. Bars in red indicate IgG from acute GBS timepoint; bars in blue indicate IgG from recovered GBS timepoint. **c** Specific glycoforms (defined in Supplementary Table 1) detected in the IgG1 subclass from acute GBS (dark red), recovered GBS (red), and purified anti-GM1 antibodies from recovered GBS donors (pink). Gal: galactose; Sia: sialic acid (Neu5Ac); AGal: without Gal; DiGal: two Gal; Fuc: fucose; Bisect: Bisecting N-glycan; Sia/Gal: Neu5Ac per Gal. Each bar represents triplicate measurements for **a** and **b**, and duplicate measurements in **c**. Bars and lines represent mean and standard deviation. * *p*-value ≤ 0.05, ** *p*-value ≤ 0.01, *** *p*-value ≤ 0.001, **** *p*-value ≤ 0.0001 determined by two-way ANOVA.

To further confirm our findings, anti-GM1 specific antibodies were purified from the three recovered GBS donors, that showed differences between their acute and recovered antibody profiles, to determine whether these overall glycoform changes were consistent with the total IgG changes observed. In each of these donors, anti-GM1 specific IgG1 antibodies were detected and confirmed to be 100% fucosylated with significantly more G1F or G2FS forms compared to total IgG from both the acute and recovered timepoints (which had detectable levels of the afucosylated antibodies NG, G1-G2) (Fig. 3b). These results provide further justification into why the G6, G8 and G19 recovered GBS donors are not relapsing.

### Anti-GM1 antibodies (Abs) cross-react to GM1-mimicking lipooligosaccharides (LOS)

We were also interested in probing whether the high titers of anti-GM1 antibodies in the recovered patients could cross-react with GM1-mimicking LOS. We surveyed 40 serum samples derived from donors including 10 healthy donors (including available household controls as indicated in Table 2), 10 enteritis donors, 10 anti-GM1 antibody positive (anti-GM1+) GBS donors and 10 anti-GM1 antibody negative (anti-GM1–) GBS donors, in western blot experiments with purified LOS from *C. jejuni* 11168, the *C. jejuni* HS:19, and engineered *Escherichia coli* CWG 308 pGM1/pCst (Supplementary Fig. 3b). *C. jejuni* 11168 and *C. jejuni* HS:19 naturally express GM1 mimics on their LOS outer core while *E. coli* CWG 308 pGM1/pCst was engineered to express GM1-mimicking LOS^32,33^. Although these strains of *C. jejuni* and *E. coli* were not isolated from our GBS patients, some anti-GM1+ donor sera still cross-reacted to these laboratory strains with GM1-mimicking LOS (Fig. 4a). Seven of our ten anti-GM1+ GBS samples exhibited cross-reactivity to at least one of the strains with varying intensity, with only six reacting to *C. jejuni* 11168 LOS. (Supplementary Fig. 3b). Additional western blot experiments were done on LOS from the *C. jejuni* 11168 mutants, *ΔneuC1* and *Δgne,* to determine whether any antibodies were generated that would recognize additional sugars beyond the ganglioside mimic (Fig. 4b). The *C. jejuni* 11168 Δ*neuC1* strain lacks N-acetylneuraminic acid (Neu5Ac) on the outer core galactose residue, while *C. jejuni* 11168 *Δgne* only expresses the inner core of LOS^34^. However, only sera from patient G99 exhibited reactivity to the inner core of the LOS (Fig. 4b). Since we confirmed that these antibodies can bind to GM1-mimicking LOS, we next examined how well these antibodies can kill *C. jejuni* 11168 in opsonophagocytosis (OPA) assays (Supplementary Fig 3c). We determined that patients with antibodies showing weak reactivity to LOS are unable to kill *C. jejuni* 11168 (G8, G66, and G19), but patients who have high reactivity to the LOS also have better OPA activities (G6, G38, G99, and H36). Further investigation of the *C. jejuni* 11168 reactivity observed for LOS isolated from proteinase K extracts led us to conclude that this reactivity was an artifact for patient G19 because two identical molecular weight bands were observed in these western blots (Fig. 4a) rather than the stepped down pattern observed in the controls (silver stain and CTB blot), and this reactivity was lost when phenol-purified LOS was examined in Fig. 4b. In contraxt, patient G8 showed a significant decrease in total IgG1 with a proportional increase in total IgG2/IgG3. These results for patient G8 and G19 likely explain the differences in *C. jejuni* OPA reactivity observed.

**Fig. 4:**
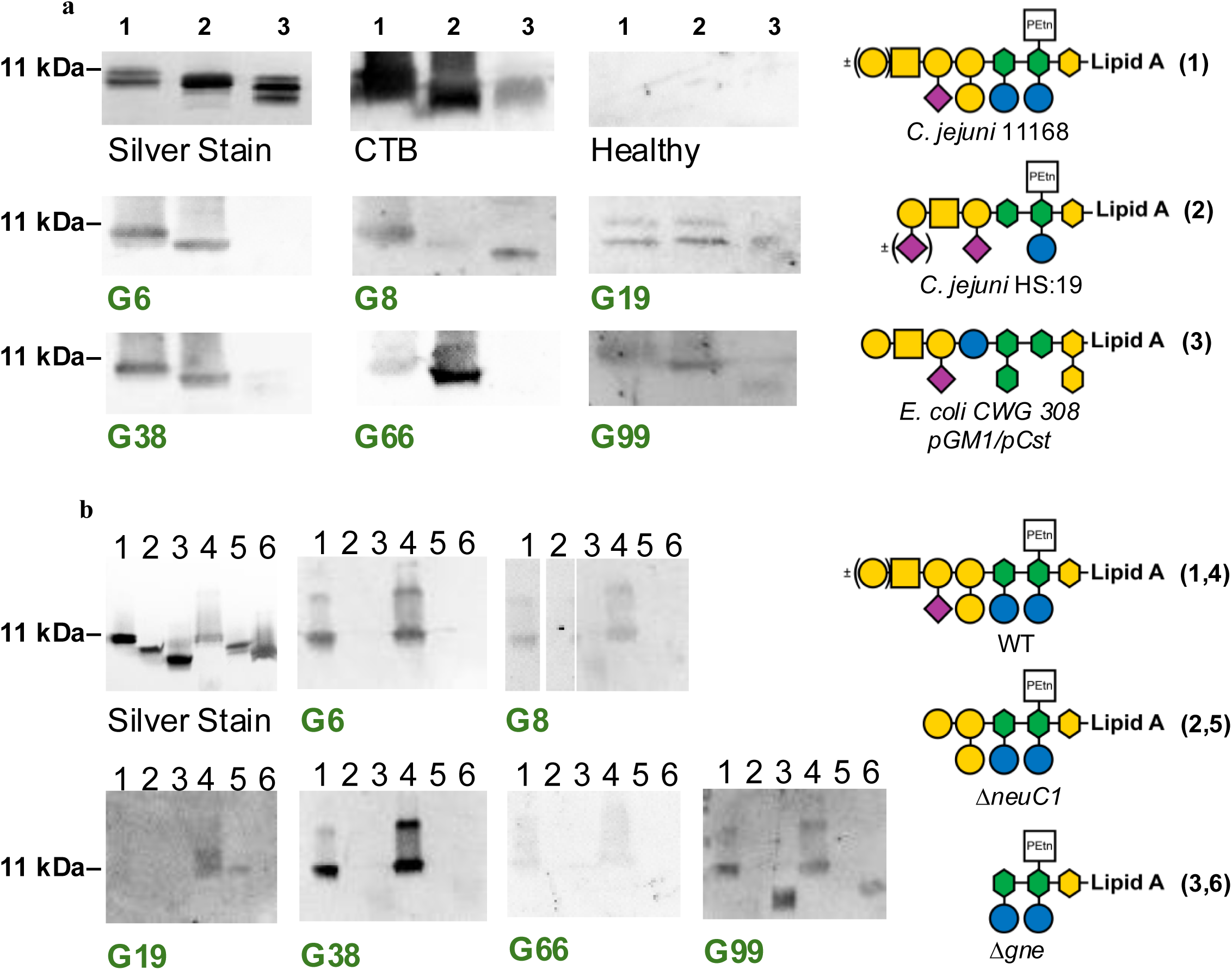
Anti-GM1 antibody positive / anti-lipooligosaccharide (LOS) antibody positive sera cross-react to GM1-mimicking bacteria. **a** Western blot analysis with various GM1 mimicking bacteria. 1. *C. jejuni* NCTC 11168 LOS 2. *C. jejuni* HS:19 serostrain LOS, and 3. *E. coli* CWG 308 *pGM1/pCst* LOS. Silver stain was used to show equal loading. Cholera toxin B subunit (CTB) was used to show reactivity to GM1-mimicking LOS. Patient ID’s in green were positive for anti-GM1 ganglioside antibodies. LOS is from proteinase K treated lysates. **b** Western blot of LOS purified via phenol preparation, lanes 1-3, and LOS isolated from proteinase K treatment, lanes 4-6. Lanes 1 and 4: *C. jejuni* NCTC 11168 wildtype (WT). Lanes 2 and 5: *C. jejuni* NCTC 11168 Δ*neuC1.* Lanes 3 and 6: *C. jejuni* NCTC 11168 Δ*gne*. Symbol nomenclature for glycans (SNFG)^43^: purple diamond, sialic acid (Neu5Ac); yellow circle, galactose (Gal); blue circle, glucose (Glc); yellow square, N-acetylgalactosamine (GalNAc); yellow hexagon, 3-deoxy-D-*manno*-octulosonic acid (KDO); green hexagon, L-*glycero*-D-*manno*-heptose (LD*man*Hep); PEtn in white square, phosphoethanolamine. The SNFG symbol in ()± indicates variation in expression of glycan.

## Discussion

In this study, we demonstrate that 10% of recovered GBS patients, from this previously studied cohort, continue to express high levels of circulating anti-GM1 antibodies in their sera equivalent to levels measured during the acute phase of their illness. This result was unexpected since the presence of anti-GM1 antibodies is the most commonly reported correlate with GBS paralysis, yet these donors no longer exhibit any signs of paralysis^14,35^. Furthermore, these titers did not correlate with time passed since GBS diagnosis or with disease severity, but did compare with high levels of anti-GM1 antibodies reported during their acute stage of illness (Supplementary Table 2). Previous studies have measured anti-ganglioside titers up to six months post GBS onset, but this is the first study to measure titers approximately ten years after initial diagnosis^2,17^. Surprisingly, 3% of healthy donors showed similar high levels of anti-GM1 antibodies, which has not been reported in previous studies. Follow-up studies examining the immune profiles, gut microbiome and IgG properties of these individuals is recommended.

Inflammation allows antibodies to pass the nerve-blood barrier^36^. To rule out the possibility that these antibodies were not causing damage to neurons from a healed nerve-blood barrier, we developed a novel GBS model that uses hPSC-derived sensory neurons to assess if these circulating antibodies could cause damage. As a proof of concept, we show complement-mediated neuropathy with commercial anti-GM1 and anti-GA1 antibodies. We then compared the pathogenicity of patient serum IgG collected at two time points of patient disease progression: acute GBS and recovered GBS. Acute phase sera from patient G19 caused more neuron destruction than even the commercial GM1-binding antibodies which could be due to observation that this sera contains antibodies to multiple gangliosides (Fig. 1b) allowing for better binding to the neuron^37,38^. After recovering, serum IgG from this patient retained the ability to bind to multiple gangliosides (Fig. 1b), but showed significantly less complement-mediated neuropathy compared to acute phase IgG, likely due to the decrease in total IgG galactosylation and increase in G2FS anti-GM1 IgG1 fucosylation^20,39^. In addition, patients who showed the greatest change in their IgG glycoforms and isotype compositions from their acute GBS and recovered GBS timepoints also showed the greatest difference in FJC-related neuropathy. These data indicate that the autoantibody pathogenicity in recovered GBS donors has decreased and no longer triggers neuropathy. Importantly, these data taken together provide evidence that IgG N-glycan modification is a primary mechanism mediating autoantibody pathogenicity in GBS.

Since these individuals no longer show any symptoms of paralysis and were not given any treatment, we hypothesized that their bodies adapted their anti-ganglioside IgG repertoire to a less inflammatory state through mechanisms of antibody selection and/or glycosyltransferase modulation. In the patient with reoccurring GBS (G38), we do not see statistically significant differences in their IgG glycome or subclass composition (Supplementary Fig. 2). However, in our other three donors, G6, G8, and G19, we see differences between their acute and recovered sera. Together, these suggest that differences in the regulation of IgG glycosylation may underlie GBS patient phenotypes, although these conclusions should be validated with additional patient data. Also, this is the first report describing anti-GM1 antibody purification from GBS patients and glycoform/subclass compositions analyses. With increased mass-spectrometry capabilities, we were also able to isolate and specifically examine the anti-GM1 IgG1 antibodies to demonstrate that increased fucosylation is likely the key factor relating to antibody pathogenesis. However, even with the larger volumes of recently collected sera, we were approaching the limit of MS detection and did not have enough sample to isolate anti-GM1 IgG1 antibodies purified from the acute sera. One limitation of this study is that it is unclear whether the anti-GM1 IgG1 antibodies have changed overtime to have less fucosylation, however we are now examining paired serum samples from US military personnel that had been banked for HIV testing and will be invaluable in addressing some of these questions.

Considering patients retain high titers of anti-GM1 antibodies years after GBS recovery, one must consider why these antibodies are present. IgG half-life is typically 21 days, although FcRn recycling may extend this^40^. These antibodies could be present due to persistent *C. jejuni* exposure. Over 30% of GBS cases are caused by an antecedent infection with *C. jejuni*^41^. Our samples were taken from the Bangladeshi population where *C. jejuni* enteritis is endemic, and up to 85% of infants are positive for *C. jejuni,* and even asymptomatic, by the age of one^42,43^. Since the frequency of *C. jejuni* infection is so high, much of the population is tolerized to the pathogen causing the rate of enteritis to decrease as they reach adulthood^10–12^. With this in mind, we hypothesized that these anti-GM1 antibodies were still in circulation due to recurrent colonization with GM1-mimicking *C. jejuni*. We therefore tested serum reactivity to LOS from ganglioside-mimicking strains of *C. jejuni*. From our 13 anti-GM1+ donors (10 GBS patients and 3 healthy controls), three patient sera (G44, G46, and G91) did not cross react with *C. jejuni* or engineered *E. coli* LOS (Supplementary Fig. 3c). Specifically, seven donors (6 GBS patients and 1 healthy donor) cross-reacted with *C. jejuni* NCTC 11168 GM1-mimicking LOS. We then performed opsonophagocytosis assays with this isolate to test the effector function of the antibodies present. We observed significant killing in half of the GBS donors with anti-GM1+ sera compared to an unrelated healthy control serum. However, further analysis of patient G19 sera demonstrated that the lack of *C. jejuni* killing was likely due to its lack of LOS recognition that we established after additional LOS purification. Sera from patient G66 had very weak reactivity to *C. jejuni* 11168 GM1-mimicking LOS, which could account for why it was also a poor opsin for this strain. In contrast, the lack of *C. jejuni* opsonophagocytosis with sera from patient G8 could be due to the significantly lower levels of IgG1 (compared to the acute phase), a subclass most associated with phagocyte recruitment due to its affinity for Fc receptors^44^. Donors with strong reactivity to *C. jejuni* 11168 LOS (G6, G38, and G99) displayed the most effective killing, most likely due to strong binding to LOS, acting as an effective opsonin for immune cells. Overall, we see that donors with strong reactivity to *C. jejuni* 11168 LOS were efficient at killing *C. jejuni*, whereas those with limited reactivity did not, suggesting one possible explanation for the retention of these autoreactive anti-GM1 antibodies.

This study found high levels of circulating anti-GM1 antibodies in a proportion of former GBS patients who currently do not exhibit signs of paralysis. The presence of anti-GM1 antibodies could in-part be due to regular immune stimulation by endemic *C. jejuni*. To avoid constant inflammatory-mediated damage, the circulating antibodies exist in a less proinflammatory through changes in N-glycosylation. For example, regular GM1 antigen exposure, could increase IgG1 fucosylation to create anti-inflammatory antibodies. In support of this finding, a previous study found repeated antigen exposure increases fucosylation for that antigen-specific antibody^45^. Furthermore, there have been several publications demonstrating that reduced fucosylation increases complement activation and antibody dependent cellular cytotoxicity leading to increased studies aimed at reducing fucose levels to enhance anti-cancer activities of antibodies^23,24,46–49^.

Our study was limited in the volume of acute sera available, and to those samples that did not lose activity after a decade of storage, so we were unable to analyze the impact of acute GBS sera in OPA assays or perform glycoform analyses of their anti-GM1 specific IgG1 antibodies. In future studies, more thorough comparisons of anti-GM1 antibody glycoforms during disease and during recovery will help to further elucidate how antibody glycosylation drives paralysis and directs recovery. With that knowledge, more effective IVIg therapies can be developed for GBS patients.

## Methods

### Blood Samples

Whole human blood was collected in GBS clinic in the National Institute of Neurosciences & Hospital and Dhaka Medical College and Hospital and icddr,b Dhaka hospital (for, *C jejuni* associated enteritis cases), in Dhaka, Bangladesh under IRB approved research protocol, PR-21060. Serum was extracted from former GBS patients (n=100), healthy donors (n=100), and *C. jejuni* associated enteritis (n=26). Collection of sera from acute GBS patients and the methods for anti-ganglioside titer calculations were previously described^28^.

### Bacterial Strains and Growth Conditions

*C. jejuni* NCTC 11168 and the *C. jejuni* HS:19 serostrain were grown on Brain Heart Infusion (BHI) agar under microaerobic conditions (5% O_2_, 10% CO_2_, and 85% N_2_). *E. coli* CWG 308 pGM1/pCst was engineered as described previously^33^. *E. coli* CWG 308 pGM1/pCst was grown on BHI with kanamycin (25 μg/mL, final concentration) and ampicillin (50 μg/mL, final concentration) and incubated at 37 °C under normal atmospheric conditions.

### Ganglioside ELISA

Healthy donors, former GBS patients, and former enteritis patients had their anti-GM1 titers analyzed in technical duplicates via ELISA (BÜHLMANN, EK-GM1-GM-U). Percent ratio was calculated as the 450 nm absorbance of samples and controls divided by the 450 nm absorbance of the calibrator. According to the manufacturer’s instructions, samples that had an absorbance 50% or higher than the calibrator were considered anti-GM1 positive. Samples that fell between 30% to 50% were considered inconclusive and below 30% were considered anti-GM1 negative. Serum from donors who were anti-GM1 positive were subsequently tested for reactivity to other anti-gangliosides (BÜHLMANN, EK-GCO-U).

### LOS Isolation

LOS was isolated as previously described^50^ with the following modifications. *C. jejuni* NCTC 11168, *C. jejuni* HS:19 serostrain, and *E. coli* CWG 308 pGM1/pCst were collected from BHI agar and set to an OD_600_ = 1 in 10 mL. Cells were pelleted and resuspended in 400 µL SDS loading buffer (100 mM Tris-Cl (pH 8.0), 2% β-mercaptoethanol, 4 % SDS, 0.2 % bromophenol blue, 0.2 % xylene cyanol, and 20 % glycerol) and boiled for 10 min at 95 °C. After cooling to room temperature, proteinase K was added to a final concentration of 500 µg/mL and cells were incubated overnight at 37°C, followed by enzyme inactivation via boiling at 95 °C for 10 min.

### LOS Silver Stain

Silver staining of bacterial LOS was performed as previously described^51^. Briefly, LOS was separated onto 12.5% SDS-PAGE gels at 110 V for 10 min and then 165 V for 65 min. The gel was then incubated with fixing solution (40% ethanol and 5% acetic acid in Milli-Q water) for 2 hours at room temperature followed by oxidizing solution (fixing solution with 0.7% periodic acid) for 15 min. After three wash steps, the gel was incubated in pre-stain (18.67 mM NaOH, 1.3% NH_4_OH in Milli-Q water) for 10 min. The silver stain (pre-stain with 39.2 mM silver nitrate) was added to the gel and stained for 10 min. The gel was then washed three times and developer (Bio-Rad) was added. Development was stopped with 5% acetic acid and the gel was imaged. All wash steps were done in Milli-Q water three times, 15 min each.

### LOS Western Blots

Samples were loaded onto 12.5% SDS-PAGE gels. Samples were run at 110 V for 10 min and then 165 V for 60 min. Samples were transferred onto nitrocellulose membrane at 100 V for 60 min. Membranes were blocked for 1 hour at room temperature in 5% bovine serum albumin (BSA) in phosphate-buffered saline with 0.05% Tween-20 (PBST). Then primary antibodies were added at the following concentrations: 1:1,000 dilution of human sera, 1:1,000 dilution of mouse-anti-GM1 (DSHB), or 1:100,000 dilution of CTB in blocking buffer to the membranes and incubated overnight at 4 °C with rocking. Membranes were rinsed quickly three times and were then washed for 5 min three times. For the secondary antibodies, goat-anti-human-HRP (Invitrogen) was added at a dilution of 1:10,000 for membranes treated with human sera. Goat-anti-mouse-HRP (Invitrogen) was added at a dilution of 1:10,000 for membranes treated with mouse-anti-GM1 and incubated at room temperature for one hour. Rabbit anti-cholera toxin (Abcam) was diluted 1:5,000, and added to the membranes treated with CTB, and incubated overnight. CTB treated membranes were washed and rinsed as stated previously, and goat-anti-rabbit-HRP (Invitrogen) was added at concentration of 1:10,000 and incubated at room temperature for one hour. Membranes were rinsed and washed as stated previously. Membranes were developed with Clarity Western ECL Substrate (BioRad) and imaged using ChemiDoc MP Gel Imaging System (BioRad).

### Antibody Isolation from Sera

IgG antibodies were isolated from sera using NAb™ Protein G Spin Columns (ThermoFisher, 89953) according to manufacturer’s instructions and eluted with Pierce™ Gentle Ag/Ab Elution Buffer, pH 6.6 (ThermoFisher, 21027). Purified IgG was concentrated to 1 mg/mL with a 10 K MWCO spin filter (ThermoFisher, 88513).

Anti-GM1 specific antibodies were purified using GM1 coated ELISA plates (BÜHLMANN, EK-GM1-GM-U) following the manufacturer’s protocol. Rather than adding the detection antibody, however, the plate was washed twice with PBST, twice with ammonium bicarbonate (ABC), and then treated with trypsin (50 ng/µL) overnight at 37 °C. Anti-GM1 specific antibodies were subsequently collected and lyophilized for MS analysis.

### Serum Killing Assays

Serum killing assays were done as previously described^52^ with the following modifications. *C. jejuni* NCTC 11168 was harvested from BHI agar, washed once with media (2 mM L-glutamine + 10 mM HEPES + 50 µM 2-mercaptoethanol + 5% heat-inactivated FBS all in RPMI 1640), and then set to an OD_600_= 0.1. Whole blood was collected in tubes pre-coated with heparin and centrifuged at 300 x *g* for 10 min so that the serum could be removed. Each well contained 40% serum from sample donor, 37.5% media, 12.5% fresh human blood with serum removed via centrifugation (500 x g for 10 min at 4°C), and 10% *C. jejuni*. Bacteria were incubated at 37 °C for 2 hours in a 5% CO_2_ incubator and mixed every 15 min with gentle agitation. After incubation, the bacteria were serially diluted in PBS and spread on BHI agar plates. After incubating for 48 hours in microaerobic conditions, colony forming units (CFU)/mL were enumerated.

### Human Pluripotent Stem Cell Culture

WA09 (also called H9) human embryonic stem cells were used in all experiments. hPSCs were cultured as described in Saito-Diaz and Zeltner^53^. Briefly, hPSCs were cultured in xeno-free culture conditions on recombinant human vitronectin (rhVTN) substrate (Gibco A31804) in Essential 8 media (Gibco A1517001). hPSC colonies were passaged every 3-4 days using EDTA solution. hPSCs were maintained below passage 60.

### Sensory Neuron (SN) Differentiation

SNs were differentiated as described in Saito-Diaz and Zeltner^53^. Briefly, on day 0, hPSCs were dissociated into single cells and seeded on rhVTN substrate at a density of 200,000 cells/cm2 in neural crest cell (NCC) differentiation media 1. NCC differentiation media 1 is comprised of Essential 6 media (Gibco A1516401) containing 10 μM SB431542 (Tocris 1614), 0.5 ng/mL recombinant human BMP4 (R&D Systems 314-BP), 300 nM CHIR99021 (Tocris 4423), and 10 μM Y-27632 (Biogems 1293823). On day 1, media is replaced with fresh NCC differentiation media 1. On day 2, media was replaced with NCC differentiation media 2 containing 0.75 μM CHIR99021, 2.5 μM DAPT (Bio-techne 2634/50), and 2.5 μM SU5402 (Biogems 2159233), while BMP4 was removed. Cultures were fed every 2 days from day 2 to day 12.

On day 12, NCCs were dissociated by 45 min accutase (Innovative Cell Technologies NC9464543) treatment and resuspended at 250,000 cells/cm^2^ on plates coated with Poly-L-ornithine (Sigma-Aldrich P3655), fibronectin (Corning 47743-654), and Laminin-1 (Cultrex 3401-010-02) in SN differentiation media. SN differentiation media contained Neurobasal medium (Gibco 21103049) with 1X N2 Supplement (Gibco 17502-048), 1X B-27 supplement without vitamin A (Gibco 12587-010), 2 mM L-glutamine (Gibco 25030-081), 20 ng/mL GDNF (R&D Systems 248-BD), 20 ng/mL BDNF (PeproTech 450-10), 25 ng/mL NGF (PeproTech 450-01), 0.6 μg/mL Laminin-1, 0.6 μg/mL Fibronectin, 0.125μM Retinoic acid (Sigma-Aldrich R2625), 1X Antibiotic-Antimycotic (Gibco 15240096), and 1 μM DAPT. On day 20, DAPT was removed from the media. SNs were fed every 2-3 days until day 30 and every 5 days thereafter. SNs were verified using immunofluorescence for the markers PRPH (Sigma-Aldrich MAB 1585) and BRN3a (Santa Cruz Biotechnology SC-377093).

### Sensory Neuron Antibody Treatment

Day 75 hPSC-derived SNs were treated for 72 hours with either no antibody, rabbit anti-GA1 antibody, purified mouse anti-GM1 antibody, purified patient antibody, or IgG from human serum (Sigma I4506). All antibody treatments were performed at 200 nM. Each group was treated either with no complement or with 2% complement-active human serum (Innovative Research ICSER10ML).

### Fluorojade C

Fluorojade-C (FJC, Avantor 76264-638) staining was performed as described in Wu *et al* 2024 with the addition of propidium iodide (PI; Sigma-Aldrich P3566, 1:20) for identification of SN cell bodies^54^. Quantification was performed by measuring the mean fluorescence intensity of FJC in SN cell bodies in ImageJ. Images were captured at 20x magnification. Any off-target differentiation was mitigated by ensuring analysis only on SNs. To this end, SN cell bodies were identified using Nissl body staining by propidium iodide^55^. Cells within SN clusters were not quantified to avoid capturing signal resulting from autofluorescence. To account for variability in background signal, the background signal intensity near each cell was also measured, then subtracted from the mean fluorescence intensity of each cell. The FJC mean fluorescence intensity for cells within each condition was then averaged per experiment. The technical replicates are defined as images taken across the same well. Only raw images were utilized for FJC quantification, and all images were taken with the same microscope settings for each image across all conditions. Fluorescence microscopy was performed on an Agilent Lionheart microscope. Image acquisition was performed using Gen5 software. Color overlays were generated in Adobe Photoshop.

### ROS Measurement

ROS levels were analyzed using the CM-H2DCFDA (Invitrogen, C6827) oxidative stress indicator. Cells were washed twice with PBS, and incubated with 10 mM CM-H2DCFDA in PBS for 45 min at 37 °C. After incubation, cells were washed twice with PBS, and fluorescent signal was measured using fluorescence microscopy imaging (AMG EVOS FL). Fluorescence intensity was quantified as described above for FJC.

### Sensory Neuron Immunofluorescence

SNs were fixed for 10 min with 4 % paraformaldehyde (Thermo Scientific AAJ19943K2) in DPBS (Corning 21-031-CM), washed three times with DPBS, permeabilized for 5 min with 0.1% Triton X-100 (Sigma X100) in DPBS, and blocked for 30 min in 0.2 % Tween-20 solution in DPBS containing 3% normal goat serum (Sigma S26-100mL), 1% BSA (Sigma A4503), and 0.01% sodium azide (Sigma S2002). SNs were then incubated with 200 nM of each antibody overnight at 4 °C in primary antibody solution comprised of DPBS containing 3% normal goat serum and 1% BSA. Then, after the primary antibody solution was removed, cells were washed 3 times with DPBS, and cells were incubated for 1 hour with a fluorophore-conjugated secondary antibody solution at room temperature containing secondary antibodies along with 5 μg/mL DAPI (Sigma D9542) as a nuclear counterstain. Secondary antibody base solution was the same as primary antibody solution. Microscopy and image processing was performed as described above for FJC.

### Mass Spectrometry Analysis

Samples for LC-MS analysis were prepared by diluting 1 µL stored serum into 50 µL of 50 mM ammonium bicarbonate (ABC). D,L-dithiothreitol (DTT) was spiked into the sample at a final concentration of 5 mM followed by incubation at 50 °C for 45 min, and then alkylated with α-iodoacetamide (IAA) to a final concentration of 13 mM, and incubated at room temperature in the dark for 45 min. Excess IAA was quenched by adding an aliquot of 5 mM DTT. The solution was combined with sequencing grade trypsin (1:25 enzyme:protein ratio) and incubated at 37 °C overnight. The digestion was halted by heating to 100°C for 5 min, diluted into LC-MS-grade water to a final concentration of 0.1 µg/µL, and filtered through 0.2 µm filters.

LC-MS analysis was carried out on an Ultimate 3000 – Orbitrap Eclipse system using an Acclaim PepMax 100 C18 nano column (ThermoScientific; 75 µm × 15 cm, 2 µm). Peptides were separated by 0-40 min 4-62.5% Buffer B (80% Acetonitrile, ACN 0.1% formic acid) in a 1 hour gradient, at a flowrate of 0.150 µL/min (Buffer A is 0.1% FA). A full scan MS1 (m/z 350-2000) was performed in the positive mode with, RF lens 60%, and Orbitrap resolution 120k. Data dependent analysis was carried out in a top 3 sec loop for precursor charges 1-6 and minimum intensity 5.0e4. Fragmentation was accomplished with stepped HCD using normalized collision energy (NCE) combinations of 20/35/45%. MS2 spectra were recorded at 30k resolution. Approximately 0.5 µg total peptide was injected for each sample.

Glycopeptide identification was carried out using Byonic software (v.5.7.33), against the conserved Fc regions of major human IgG with peak integration carried out in FreeStyle 1.8 SP2 QF1. MS1 deconvoluted chromatograms were fitted by Xtract algorithm considering 1-6 charges. Deconvoluted XIC was drawn with 10 ppm parent mass error which was used to report relative abundance in each sample. Glycan traits were used for unified comparison among samples to compensate for detection fluctuations between samples and intrinsic interdependent of glycosylation within samples.

### Statistics

All statistical tests were calculated using GraphPad Prism version 10. For the anti-GM1 ELISA, Kruskal-Wallis test was used. Serum killing assays were tested via one-way ANOVA. Significance for LC-MS glycosylation profiles were done with two-way ANOVA. FJC and ROS analyses were tested with one-way ANOVA. P values less than 0.05 were considered significant.

## Acknowledgements

We thank Dr. Balazs Rada lab for providing blood for use in the OPA experiments supported by the National Institutes of Health (NIH) under award number 5R01HL136707 and R21AI147097; and also thank the healthy volunteers for their blood donations, the staff of the UGA Health Center, and the UGA Clinical and Translational Research Unit for drawing blood used in the OPA studies. We thank Drs. Linda Mansfield, Harald Nothaft and Leandre Glendenning for their helpful discussions. We also thank Drs. James and Adrienne Paton for providing us with their *E. coli* CWG 308 wildtype and pGM1/pCst strains.

## Author Contributions

C.M.S., Z.I., and N.Z. conceived the project. N.P., S.H., and I.J. and Z.I. enrolled previous GBS patients and obtained their medical history. A.M.R performed, analyzed, and interpreted the ELISA and OPA experiments. A.M.R purified antibodies for neuron and MS experiments. X.Y. and S.A.H performed mass spectrometry experiments, analyzed the data, and interpreted the data with A.M.R. J.M performed neuron experiments and data analysis. A.M.R. wrote the manuscript and it was edited by all the co-authors. C.M.S., P.A., Z.I., and N.Z. supervised the project.

## References

1. McGrogan, A., Madle, G. C., Seaman, H. E. & De Vries, C. S. The Epidemiology of Guillain-Barré Syndrome Worldwide. A Systematic Literature Review. Neuroepidemiology 32, 150–163 (2009).

2. Thomma, R. C. M., Fokke, C., Walgaard, C., Vermeulen-De Jongh, D. M. C., Tio-Gillen, A., van Rijs, W., van Doorn, P. A., Huizinga, R. & Jacobs, B. C. High and Persistent Anti-GM1 Antibody Titers Are Associated With Poor Clinical Recovery in Guillain-Barré Syndrome. Neurol. Neuroimmunol. Neuroinflamm. 10, e200107 (2023).

3. Islam, Z., Gilbert, M., Mohammad, Q. D., Klaij, K., Li, J., van Rijs, W., Tio-Gillen, A. P., Talukder, K. A., Willison, H. J., van Belkum, A., Endtz, H. P. & Jacobs, B. C. Guillain-Barré syndrome-related *Campylobacter jejuni* in Bangladesh: Ganglioside mimicry and cross-reactive antibodies. PLoS One 7, (2012).

4. World Health Organization. Campylobacter. (2020).

5. Baker, M. G., Kvalsvig, A., Zhang, J., Lake, R., Sears, A. & Wilson, N. Declining Guillain-Barré Syndrome after Campylobacteriosis Control, New Zealand, 1988–2010. Emerg. Infect. Dis. 18, 226–233 (2012).

6. Yuki, N., Susuki, K., Koga, M., Nishimoto, Y., Odaka, M., Hirata, K., Taguchi, K., Miyatake, T., Furukawa, K., Kobata, T. & Yamada, M. Carbohydrate mimicry between human ganglioside GM1 and *Campylobacter jejuni* lipooligosaccharide causes Guillain-Barré syndrome. Proc. Natl. Acad. Sci. U. S. A. 101, 11404–11409 (2004).

7. Ogawa-Goto, K., Funamoto, N., Abe, T. & Nagashima, K. Different ceramide compositions of gangliosides between human motor and sensory nerves. J. Neurochem. 55, 1486–1493 (1990).

8. Griffin, J. W., Li, C. Y., Ho, T. W., Tian, M., Gao, C. Y., Xue, P., Mishu, B., Cornblath, D. R., Macko, C., McKhann, G. M. & Asbury, A. K. Pathology of the motor-sensory axonal guillain-barré syndrome. Ann. Neurol. 39, 17–28 (1996).

9. Godschalk, P. C. R., Kuijf, M. L., Li, J., St. Michael, F., Ang, C. W., Jacobs, B. C., Karwaski, M. F., Brochu, D., Moterassed, A., Endtz, H. P., Van Belkum, A. & Gilbert, M. Structural characterization of *Campylobacter jejuni* lipooligosaccharide outer cores associated with Guillain-Barré and Miller Fisher syndromes. Infect. Immun. 75, 1245–1254 (2007).

10. Susuki, K., Rasband, M. N., Tohyama, K., Koibuchi, K., Okamoto, S., Funakoshi, K., Hirata, K., Baba, H. & Yuki, N. Anti-GM1 antibodies cause complement-mediated disruption of sodium channel clusters in peripheral motor nerve fibers. J. Neurosci. 27, 3956–3967 (2007).

11. Paparounas, K., O’Hanlon, G. M., O’Leary, C. P., Rowan, E. G. & Willison, H. J. Anti-ganglioside antibodies can bind peripheral nerve nodes of Ranvier and activate the complement cascade without inducing acute conduction block in vitro. Brain 122, 807–816 (1999).

12. Rodella, U., Scorzeto, M., Duregotti, E., Negro, S., Dickinson, B. C., Chang, C. J., Yuki, N., Rigoni, M. & Montecucco, C. An animal model of Miller Fisher syndrome: Mitochondrial hydrogen peroxide is produced by the autoimmune attack of nerve terminals and activates Schwann cells. Neurobiol. Dis. 96, 95–104 (2016).

13. Nobile-Orazio, E., Carpo, M., Meucci, N., Grassi, M. P., Capitani, E., Sciacco, M., Mangoni, A. & Scarlato, G. Guillain-Barré syndrome associated with high titers of anti-GM1 antibodies. J. Neurol. Sci. 109, 200–206 (1992).

14. Kuwabara, S., Yuki, N., Koga, M., Hattori, T., Matsuura, D., Miyake, M. & Noda, M. IgG anti-GM1 antibody is associated with reversible conduction failure and axonal degeneration in Guillain-Barré syndrome. Ann. Neurol. 44, 202–208 (1998).

15. Ogawara, K., Kuwabara, S., Koga, M., Mori, M., Yuki, N. & Hattori, T. Anti-GM1b IgG antibody is associated with acute motor axonal neuropathy and *Campylobacter jejuni* infection. J. Neurol. Sci. 210, 41–45 (2003).

16. Van Sorge, N. M., Van den Berg, L. H., Geleijns, K., Van Strijp, J. A., Jacobs, B. C., Van Doorn, P. A., Wokke, J. H. J., Van de Winkel, J. G. J., Leusen, J. H. W. & Van der Pol, W. L. Anti-GM1 IgG antibodies induce leukocyte effector functions via Fcγ receptors. Ann. Neurol. 53, 570–579 (2003).

17. Press, R., Matá, S., Lolli, F., Zhu, J., Andersson, T. & Link, H. Temporal profile of anti-ganglioside antibodies and their relation to clinical parameters and treatment in Guillain-Barré syndrome. J. Neurol. Sci. 190, 41–47 (2001).

18. Intravenous immunoglobulin for Guillain-Barré syndrome - Hughes, RAC - 2014 | Cochrane Library. https://www.cochranelibrary.com/cdsr/doi/10.1002/14651858.CD002063.pub6/full.

19. Buhre, J. S., Becker, M. & Ehlers, M. IgG subclass and Fc glycosylation shifts are linked to the transition from pre- to inflammatory autoimmune conditions. Front. Immunol. 13, (2022).

20. Wei, B. et al. Fc galactosylation follows consecutive reaction kinetics and enhances immunoglobulin G hexamerization for complement activation. MAbs 13, (2021).

21. Quast, I., Keller, C. W., Maurer, M. A., Giddens, J. P., Tackenberg, B., Wang, L. X., Münz, C., Nimmerjahn, F., Dalakas, M. C. & Lünemann, J. D. Sialylation of IgG Fc domain impairs complement-dependent cytotoxicity. Journal of Clinical Investigation 125, 4160– 4170 (2015).

22. Fokkink, W. J. R., Selman, M. H. J., Dortland, J. R., Durmuş, B., Kuitwaard, K., Huizinga, R., Van Rijs, W., Tio-Gillen, A. P., Van Doorn, P. A., Deelder, A. M., Wuhrer, M. & Jacobs, B. C. IgG Fc N-glycosylation in Guillain-Barré syndrome treated with immunoglobulins. J. Proteome Res. 13, 1722–1730 (2014).

23. Van Coillie, J., Schulz, M. A., Bentlage, A. E. H., de Haan, N., Ye, Z., Geerdes, D. M., van Esch, W. J. E., Hafkenscheid, L., Miller, R. L., Narimatsu, Y., Vakhrushev, S. Y., Yang, Z., Vidarsson, G. & Clausen, H. Role of N-Glycosylation in FcγRIIIa interaction with IgG. Front. Immunol. 13, 987151 (2022).

24. Gasdaska, J. R., Sherwood, S., Regan, J. T. & Dickey, L. F. An afucosylated anti-CD20 monoclonal antibody with greater antibody-dependent cellular cytotoxicity and B-cell depletion and lower complement-dependent cytotoxicity than rituximab. Mol. Immunol. 50, 134–141 (2012).

25. Martin, T. C. et al. Decreased Immunoglobulin G Core Fucosylation, A Player in Antibody-dependent Cell-mediated Cytotoxicity, is Associated with Autoimmune Thyroid Diseases. Molecular & Cellular Proteomics 19, 774–792 (2020).

26. Ząbczyńska, M., Link-Lenczowski, P. & Pocheć, E. Glycosylation in Autoimmune Diseases. Adv. Exp. Med. Biol. 1325, 205–218 (2021).

27. Vučkovïc, F. et al. Association of systemic lupus erythematosus with decreased immunosuppressive potential of the IgG glycome. Arthritis and Rheumatology 67, 2978– 2989 (2015).

28. Halstead, S. K., Kalna, G., Islam, M. B., Jahan, I., Mohammad, Q. D., Jacobs, B. C., Endtz, H. P., Islam, Z. & Willison, H. J. Microarray screening of Guillain-Barré syndrome sera for antibodies to glycolipid complexes. Neurol. Neuroimmunol. Neuroinflamm. 3, 1–9 (2016).

29. Huang, C. W., Rust, N. C., Wu, H. F., Yin, A., Zeltner, N., Yin, H. & Hart, G. W. Low glucose induced Alzheimer’s disease-like biochemical changes in human induced pluripotent stem cell-derived neurons is due to dysregulated O-GlcNAcylation. Alzheimer’s and Dementia 19, 4872–4885 (2023).

30. Ikenari, T., Kurata, H., Satoh, T., Hata, Y. & Mori, T. Evaluation of Fluoro-Jade C Staining: Specificity and Application to Damaged Immature Neuronal Cells in the Normal and Injured Mouse Brain. Neuroscience 425, 146–156 (2020).

31. Teng, L., Fan, L., Peng, Y., He, X., Chen, H., Duan, H., Yang, F., Lin, D., Lin, Z., Li, H. & Shao, B. Carnosic Acid Mitigates Early Brain Injury After Subarachnoid Hemorrhage: Possible Involvement of the SIRT1/p66shc Signaling Pathway. Front. Neurosci. 13, 417646 (2019).

32. Yu, R. K., Usuki, S. & Ariga, T. Ganglioside molecular mimicry and its pathological roles in Guillain-Barré syndrome and related diseases. Infect. Immun. 74, 6517–6527 (2006).

33. Focareta, A., Paton, J. C., Morona, R., Cook, J. & Paton, A. W. A Recombinant Probiotic for Treatment and Prevention of Cholera. Gastroenterology 130, 1688–1695 (2006).

34. Guerry, P., Szymanski, C. M., Prendergast, M. M., Hickey, T. E., Ewing, C. P., Pattarini, D. L. & Moran, A. P. Phase variation of *Campylobacter jejuni* 81-176 lipooligosaccharide affects ganglioside mimicry and invasiveness in vitro. Infect. Immun. 70, 787–793 (2002).

35. Campbell, C. I., Mcgonigal, R., Barrie, J. A., Delaere, J., Bracke, L., Cunningham, M. E., Yao, D., Delahaye, T., Van De Walle, I. & Willison, H. J. Complement inhibition prevents glial nodal membrane injury in a GM1 antibody-mediated mouse model. Brain Commun. 4, (2022).

36. Shimizu, F., Sato, R., Mizukami, Y., Watanabe, K., Maeda, T., Kanda, T., Matsui, N., Misawa, S., Izumi, Y., Kuwabara, S. & Nakamori, M. Blood–Nerve Barrier Breakdown Induced by Immunoglobulin G in Typical and Multifocal Chronic Inflammatory Demyelinating Polyneuropathy and Multifocal Motor Neuropathy. Int. J. Mol. Sci. 27, 1088 (2026).

37. Ogino, M., Orazio, N. & Latov, N. IgG anti-GM1 antibodies from patients with acute motor neuropathy are predominantly of the IgG1 and IgG3 subclasses. J. Neuroimmunol. 58, 77– 80 (1995).

38. Goodfellow, J. A., Bowes, T., Sheikh, K., Odaka, M., Halstead, S. K., Humphreys, P. D., Wagner, E. R., Yuki, N., Furukawa, K., Furukawa, K., Plomp, J. J. & Willison, H. J. Overexpression of GD1a ganglioside sensitizes motor nerve terminals to anti-GD1a antibody-mediated injury in a model of acute motor axonal neuropathy. Journal of Neuroscience 25, 1620–1628 (2005).

39. Peschke, B., Keller, C. W., Weber, P., Quast, I. & Lünemann, J. D. Fc-Galactosylation of Human Immunoglobulin Gamma Isotypes Improves C1q Binding and Enhances Complement-Dependent Cytotoxicity. Front. Immunol. 8, (2017).

40. Bas, M., Terrier, A., Jacque, E., Dehenne, A., Pochet-Béghin, V., Beghin, C., Dezetter, A.-S., Dupont, G., Engrand, A., Beaufils, B., Mondon, P., Fournier, N., de Romeuf, C., Jorieux, S., Fontayne, A., Mars, L. T. & Monnet, C. Fc Sialylation Prolongs Serum Half-Life of Therapeutic Antibodies. The Journal of Immunology 202, 1582–1594 (2019).

41. Allos, B. M. Association between *Campylobacter* infection and Guillain-Barré syndrome. Journal of Infectious Diseases 176, 125–128 (1997).

42. Miller, M. et al. The MAL-ED study: a multinational and multidisciplinary approach to understand the relationship between enteric pathogens, malnutrition, gut physiology, physical growth, cognitive development, and immune responses in infants and children up to 2 years of age in resource-poor environments. Clin. Infect. Dis. 59 Suppl 4, S193–S206 (2014).

43. Amour, C. et al. Epidemiology and Impact of *Campylobacter* Infection in Children in 8 Low-Resource Settings: Results From the MAL-ED Study. Clin. Infect. Dis. 63, 1171–1179 (2016).

44. Vidarsson, G., Dekkers, G. & Rispens, T. IgG subclasses and allotypes: From structure to effector functions. Front. Immunol. 5, 1–17 (2014).

45. Guo, N., Liu, Y., Masuda, Y., Kawagoe, M., Ueno, Y., Kameda, T. & Sugiyama, T. Repeated immunization induces the increase in fucose content on antigen-specific IgG N-linked oligosaccharides. Clin. Biochem. 38, 149–153 (2005).

46. Shields, R. L., Lai, J., Keck, R., O’Connell, L. Y., Hong, K., Gloria Meng, Y., Weikert, S. H. A. & Presta, L. G. Lack of fucose on human IgG1 N-linked oligosaccharide improves binding to human FcγRIII and antibody-dependent cellular toxicity. Journal of Biological Chemistry 277, 26733–26740 (2002).

47. Chenoweth, A. M. et al. An Fc-Engineered Glycomodified Antibody Supports Proinflammatory Activation of Immune Effector Cells and Restricts Progression of Breast Cancer. Cancer Res. 85, 4521–4540 (2025).

48. Mössner, E. et al. Increasing the efficacy of CD20 antibody therapy through the engineering of a new type II anti-CD20 antibody with enhanced direct and immune effector cell-mediated B-cell cytotoxicity. Blood 115, 4393–4402 (2010).

49. Matsumoto, Y., Jia, N., Heimburg-Molinaro, J. & Cummings, R. D. Targeting Tn-positive tumors with an afucosylated recombinant anti-Tn IgG. Sci. Rep. 13, 5027 (2023).

50. Westphal, O. & Jann, K. Bacterial lipopolysaccharides extraction with phenol-water and further applications of the procedure. Methods Carbohydrate Chem 5, 83–91 (1965).

51. Patry, R. T., Stahl, M., Perez-Munoz, M. E., Nothaft, H., Wenzel, C. Q., Sacher, J. C., Coros, C., Walter, J., Vallance, B. A. & Szymanski, C. M. Bacterial AB 5 toxins inhibit the growth of gut bacteria by targeting ganglioside-like glycoconjugates. Nat. Commun. 10, (2019).

52. Nothaft, H., Perez-Muñoz, M. E., Yang, T., Murugan, A. V. M., Miller, M., Kolarich, D., Plastow, G. S., Walter, J. & Szymanski, C. M. Improving Chicken Responses to Glycoconjugate Vaccination Against *Campylobacter jejuni*. Front. Microbiol. 12, (2021).

53. Saito-Diaz, K. & Zeltner, N. A protocol to differentiate nociceptors, mechanoreceptors, and proprioceptors from human pluripotent stem cells. STAR Protoc. 3, 101187 (2022).

54. Wu, H. F., Saito-Diaz, K., Huang, C. W., McAlpine, J. L., Seo, D. E., Magruder, D. S., Ishan, M., Bergeron, H. C., Delaney, W. H., Santori, F. R., Krishnaswamy, S., Hart, G. W., Chen, Y. W., Hogan, R. J., Liu, H. X., Ivanova, N. B. & Zeltner, N. Parasympathetic neurons derived from human pluripotent stem cells model human diseases and development. Cell Stem Cell 31, 734–753.e8 (2024).

55. Niu, J., Li, C., Wu, H., Feng, X., Su, Q., Li, S., Zhang, L., Yew, D. T. W., Cho, E. Y. P. & Sha, O. Propidium iodide (PI) stains Nissl bodies and may serve as a quick marker for total neuronal cell count. Acta Histochem. 117, 182–187 (2015).

56. Wuhrer, M., Stam, J. C., Van De Geijn, F. E., Koeleman, C. A. M., Verrips, C. T., Dolhain, R. J. E. M., Hokke, C. H. & Deelder, A. M. Glycosylation profiling of immunoglobulin G (IgG) subclasses from human serum. Proteomics 7, 4070–4081 (2007).

57. Varki, A. et al. Symbol Nomenclature for Graphical Representations of Glycans. Glycobiology 25, 1323–1324 (2015).

