## Supplemental File for "Autoantibodies from recovered Guillain-Barré syndrome patients exhibit altered effector functions that prevent subsequent neuron degeneration"

**Supplementary Table 1:** Antibody IgG glycoforms included in each category, as shown in Fig. 3c.

| Glycosylation | Glycoforms |
|---|---|
| Galactosylation | 0.5*(G1+G1N+G1S+G1NS+G1FN+G1F+G1FS+G1FNS) +G2+G2N+G2S+G2NS+G2F+G2FN+G2FS+G2FNS |
| Agalactosylation | G0+G0N+G0F+G0FN |
| Digalactosylation | G2+G2N+G2S+G2NS+G2F+G2FN+G2FS+G2FNS |
| Fucosylation | G0F+G1F+G2F+G0FN+G1FN+G2FN+G1FS+ G2FS+G1FNS+G2FNS |
| Bisection | G0N+G1N+G2N+G1NS+G2NS+G0FN+G1FN+G2GN+G1F NS+G2FNS |
| Sialylation | G1S+G2S+G1NS+G2NS+G1FS+G2FS+G1FNS+G2FNS |

**Supplementary Table 2:** Anti-GM1 IgG antibody titer during acute phase of initial GBS diagnosis. Titer values represent the inverse serum dilution required for 50% maximal antibody binding.

| Patient ID | anti-GM1 titer at time of diagnosis |
|---|---|
| G3 | 2147.51 |
| G6 | 50575.86 |
| G8 | 50437.09 |
| G10 | 3784.92 |
| G11 | 4489.90 |
| G12 | 3594.59 |
| G13 | 2138.15 |
| G16 | 2217.14 |
| G19 | 3234.59 |
| G20 | 1320.30 |
| G23 | 2625.79 |
| G24 | 2635.13 |
| G25 | 36977.22 |
| G26 | 12615.65 |
| G28 | 18265.70 |
| G29 | 230874.37 |
| G30 | 49603.64 |
| G37 | 2516.89 |
| G38 | 130389.63 |
| G39 | 8333.22 |
| G45 | 26159.94 |
| G58 | 16117.50 |

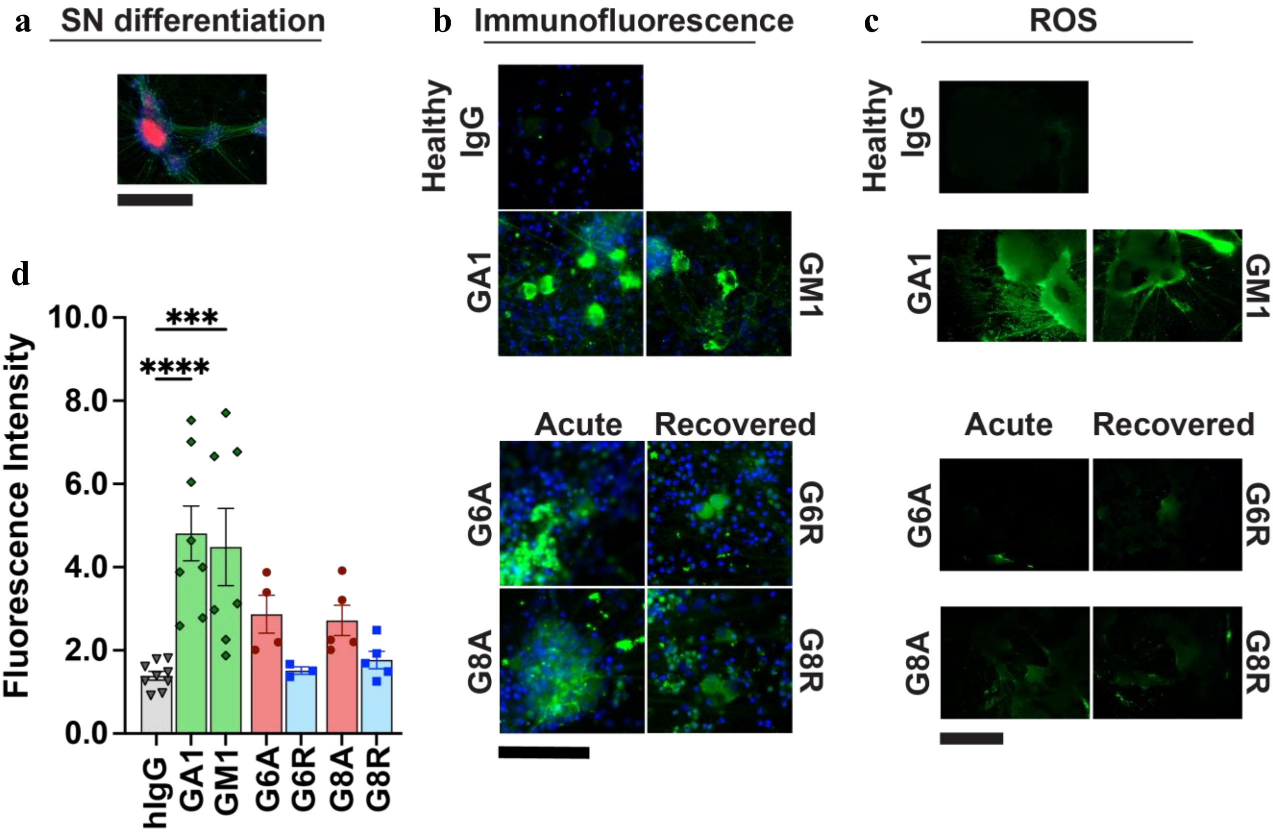


**Supplementary Fig. 1: Sensory neuron (SN) degeneration after treatment with IgG. a** SN identity was validated with immunofluorescence (IF) for the markers PRPH (green) and BRN3A (red). Scale bar: 1000 µm. **b** IF of commercial and patient antibody (Abs) binding to SN (green) and nuclear counterstain DAPI (blue). Scale bar: 100 µm. **c** SN stress and degeneration was characterized using CM-H2DCFDA (green), a reagent that detects reactive oxidative stress (ROS). Scale bar: 2000 µm. **d** Quantification of ROS fluorescence intensity. Patient samples (red, acute; blue, recovered) compared to healthy human IgG (hIgG) (grey) by one-way ANOVA. Commercial anti-GM1 (green) and anti-asialoGM1 (GA1) (green) included to show damage induced by specific anti-ganglioside Abs. Each dot represents average of technical duplicates from one donor. Lines and bars represent mean and standard error of mean. *** $p$-value $\leq 0.001$, **** $p$-value $\leq 0.0001$ determined by one-way ANOVA.

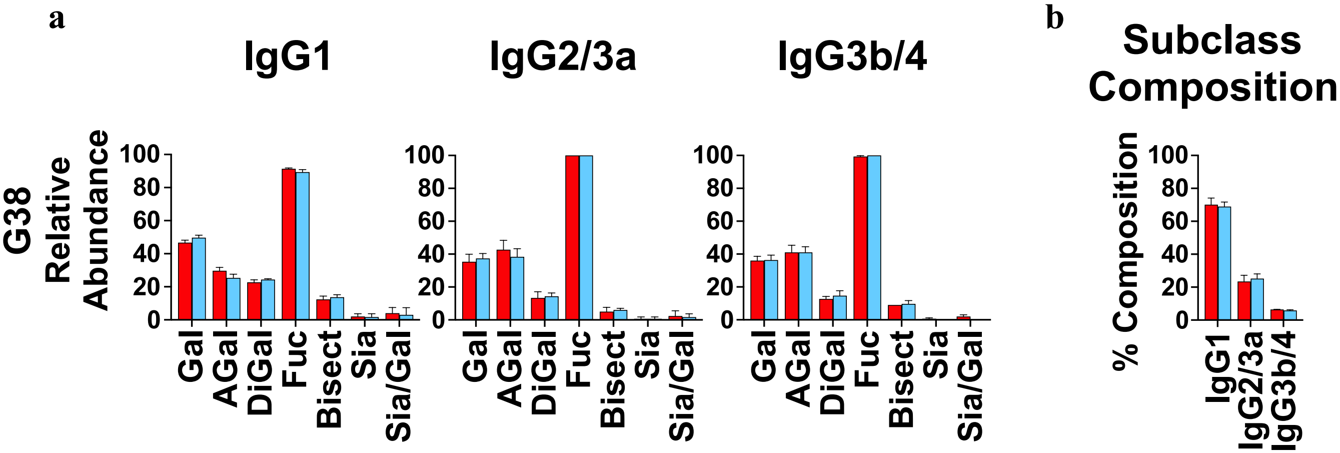


**Supplementary Fig. 2: Liquid chromatography-mass spectrometry analysis of total IgG from patient with recurring GBS. a** Glycosylation of the Fc portion of IgG separated by subclasses IgG1, IgG2/3a, and IgG3b/4. Relative abundance refers to normalized peak area detected for each glycoform. **b** Percent composition of each IgG subclass. Bars in red indicate IgG from acute GBS timepoint; bars in blue indicate IgG from recovered GBS timepoint. Each bar represents an average of triplicate measurements. Gal: galactose; Sia: sialic acid (Neu5Ac); AGal: without Gal; DiGal: two Gal; Fuc: fucose; Bisect: Bisecting N-glycan; Sia/Gal: Neu5Ac per Gal.

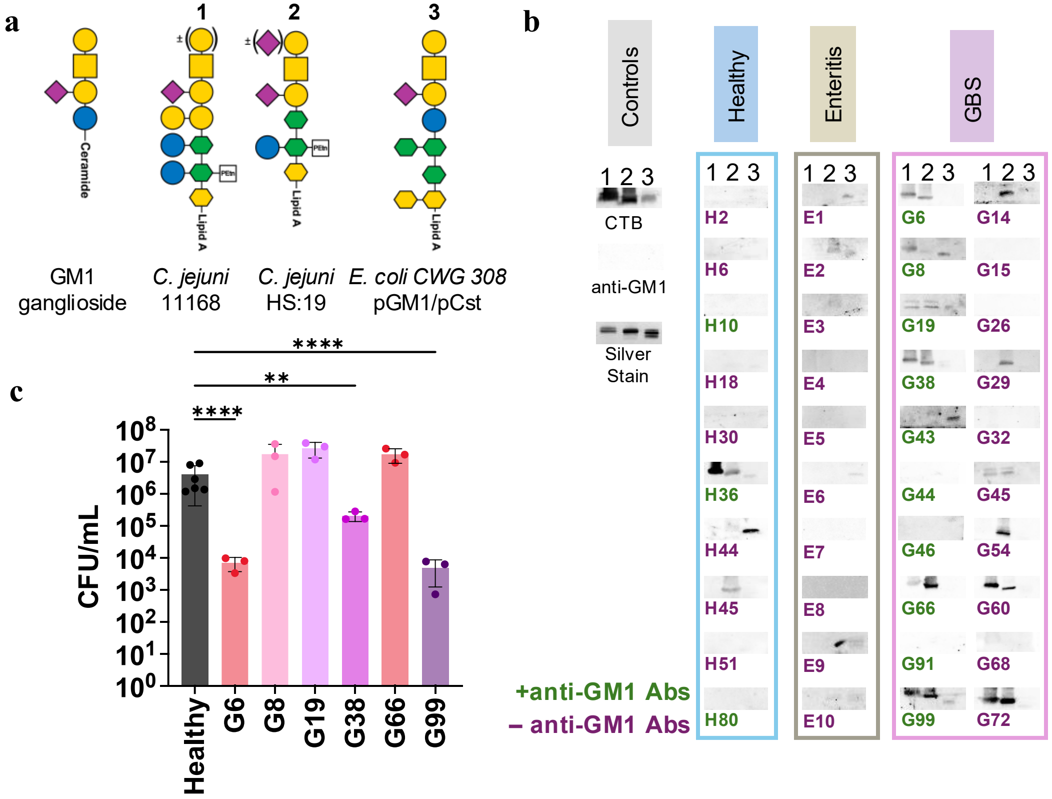


**Supplementary Fig. 3: Circulating anti-GM1 antibodies (Abs) cross react with GM1-ganglioside mimicking lipooligosaccharides (LOS). a** Schematic of GM1-ganglioside mimicking LOS structures used in this study compared with human GM1 ganglioside. LOS from: 1. *C. jejuni (Cj)* NCTC 11168, 2. *Cj* HS:19 serostrain, and 3. *E. coli* CWG 308 pGM1/pCst. **b** Western blot with healthy, *Cj* enteritis, and GBS donor sera with GM1-mimicking LOS from *Cj* 11168, *Cj* HS:19, and *E. coli* CWG 308. Controls: cholera toxin B subunit (CTB) binds GM1-mimicking structures and silver stain shows equal loading of LOS. Donor green IDs were positive for anti-GM1 ganglioside Abs (+GM1 Abs), and red IDs were negative (–GM1 Abs). **c** *Cj* 11168 colony forming units (CFU) per mL after treatment with anti-GM1+/anti-LOS+ sera and fresh whole blood. Healthy sera from donor outside this study cohort. Symbol nomenclature for glycans (SNFG)[56]: purple diamond, Neu5Ac; yellow circle, Gal; blue circle, Glc; yellow square, GalNAc; yellow hexagon, KDO; green hexagon, LD*man*Hep; PEtn in white square, phosphoethanolamine. The SNFG symbol in ()± indicates variation in glycan expression.
